# White matter hyperintensity volume and early-onset post-stroke depression in Thai older patients

**DOI:** 10.1101/2021.01.29.428768

**Authors:** Chaichana Jaroonpipatkul, Jaruwan Onwanna, Chavit Tunvirachaisakul, Nutchawan Jittapiromsak, Yothin Rakvongthai, Aurauma Chutinet, Thitiporn Supasitthumrong

## Abstract

**Objective:** Post-stroke depression (PSD) is one of the most frequent psychiatric symptoms after a stroke event. The role of white matter hyperintensities (WMHs) associated with PSD in older patients remains unclear. This study aimed to examine the volume and location of white matter microstructure abnormalities among older patients with early-onset PSD.

**Methods:** Older (≥55 years) patients with acute cerebral infarction and hospitalized in King Chulalongkorn Memorial Hospital’s stroke unit from October 2019 to September 2020 were recruited. Participants were assessed with the Montgomery-Åsberg Depression Rating Scale (MADRS) within three months after the onset of stroke. All patients had MRI scans. The brain images were segmented into four regions via left/right, frontal/dorsal plains. Two WMHs volume detections (visual rating vs. semi-automated WMHs volumetric detection) were employed on the fluid-attenuated inversion recovery images (FLAIR) for each segment. The study then investigated the association between WMHs volume and MADRS score with regression analysis.

**Results:** The study included twenty-nine patients with acute stroke. Total WMHs volume and segmented regions were not statistically associated with the MADRS score. However, there was a trend in different WHMs volume of the left anterior segment between depressed and non-depressed groups (t-test 2.058, p = 0.055). Further, demographic and clinical data showed no association with depressive symptoms.

**Conclusion:** The volume of WHMs might not contribute to the development of early-onset PSD in older patients. This study showed a potential of a quantitative MRI analysis in clinical practice. Further investigation with a larger group of patients is needed.

## Introduction

Stroke is one of the leading neurological causes of mortality worldwide,[1] and has deleterious consequences after the early and delayed onset of stroke. According to the World Stroke Organization, there are over 13.7 million new strokes each year, and five and a halfmillion people die from stroke annually worldwide.[2] Post-stroke depression (PSD) is one of the most common mood disorders following a stroke, with estimated prevalence rates between 6% and 79%,[3] particularly in the first three months after stroke. Stroke survivors will have suffered from PSD if left under-recognized leading to functional disability, poorer rehabilitation outcomes, and increased morbidity and mortality than non-depressed patients.[4] The mortality rate in Post-stroke depressed patients has been reported higher than in non-depressed. Moreover, PSD has a negative effect on rehabilitation outcomes and functional recovery.[5] Therefore, early detection of PSD is the one priority.[6] All depressive symptoms should be evaluated to deploy early intervention and management indicated since the time of patients’ hospitalization.

There are two onset periods of PSD. An early-onset refers to depression emerging within three months after the incidence of stroke, and a late-onset refers to depression occurring three months after stroke.[7] According to DSM-5, depressive symptoms are described by a depressed mood and diminished interest or pleasure in participating in activities.[8] Associated symptoms are necessary for diagnosing depressive disorder, including feelings of worthlessness, excessive or inappropriate guilt, reduced ability to think or concentrate, indecisiveness, and recurrent thoughts of death or recurrent suicidal ideation. The depressive symptoms might be coupled with somatic symptoms, such as a significant weight loss, changing (decrease or increase) in appetite, slowing down of thought, reducing physical movement or psychomotor agitation, fatigue or loss of energy, and sleep disturbance.[8] The clinical manifestation of depression after stroke does not differ from depression in other patient populations.[9] The symptoms profile of depression in stroke patients is similar to depressed patients in other illnesses seen in general practice. However, appetite disturbance and fatigue might be critical symptoms to suggest PSD during acute stroke. [10]

Evidence suggests that an association between white matter lesions and depression [11, 12]. White matter lesions, also known as white matter hyperintensities(WMHs) or leukoaraiosis, refer to the punctate or patchy changes mainly in the periventricular and subcortical white matter of the brain.[13] The limbic-cortical-striatal-pallidal-thalamic (LCSPT) circuit,[14] seen as extensively interconnected white matter fiber, is associated with affective disorder and negative emotion. [15] Changes in the white matter from an acute cerebrovascular accident could lead to the interruption of the LCSPT circuit and might contribute to the emergence of post-stroke depression. Although many lesions in various brain areas are associated with PSD, WMHs volume showed the most robust relationship with depression [16].

Various brain imaging studies examined the lesion locations of stroke linked to PSD, such as frontal lobe [17, 18], temporal lobe[19], occipital-parietal lobe[20], subcortical[21], basal ganglia[22], thalamus[23] and insular cortex.[24] Previous studies reported that left anterior brain injury was associated with depressive mood, while the right side lesion showed the reverse trend such as overly cheerful, apathy and anosognosia.[25, 26] However, another study suggested that the right hemisphere was also related to PSD.[27] Although it is still inconclusive, previous studies have demonstrated that patients with lesions in the frontal lobe had a higher risk of PSD and have provided some pathophysiological underpinnings for PSD.[21] For white matter lesions, previous studies found that the pathological changes in the white matter microstructure in the frontal lobe, temporal lobe, and genu of corpus callosum might underlie the mechanism of PSD.[27] Williamson et al., 2010 [26], using MRI and calculated fractional anisotropy, found that frontal and parietal white matter lesions were more strongly and consistently related to mood scores than other regions. However, it remains unclear what relationship exists between lesions location and the volume of white matter hyperintensities in the emergence of PSD, particularly in the early onset type.

Magnetic resonance imaging (MRI) provides a much higher image resolution than computed tomography (CT). MRI is also more sensitive in identifying the site and extent of the infarct and in revealing white matter lesions [28]. However, previous studies on PSD employed CT, and there has been a lack of MRI studies in this specific field of investigation.[29] Therefore, our study aimed to use MRI to estimate white matter lesions’ volume and location to determine any association with depression after stroke. We also used diffusion-weighted magnetic resonance imaging (DWI) to measure acute vascular lesions. DWI shows the infarcted core in ischemic stroke patients, which indicates the actual minimum extension of infarction and has an excellent interrater agreement. Additionally, DWI has substantially better sensitivity and accuracy than CT.[30]

This study aimed to identify if WMHs volume and location correlated with early-onset PSD among older patients in King Chulalongkorn Memorial Hospital. Any study findings might help clinicians predict and deliver proper management for patients who might have PSD.

## Methods

### Subjects

Study approval was obtained from the Institutional Review Board of the Faculty of Medicine, Chulalongkorn University, Bangkok, Thailand, (IRB no.145/63), which is in compliance with the International Guideline for Human Research protection, as required by the Declaration of Helsinki, The Belmont Report, CIOMS Guideline, and International Conference on Harmonization in Good Clinical Practice (ICH-GCP). Patients with acute cerebral infarction hospitalized at the King Chulalongkorn Memorial hospital’s stroke unit from October 2019 to September 2020 were recruited. Inclusion criteria were acute ischemic stroke patients age 55 years or older who were diagnosed with focal neurological signs or symptoms thought to be of vascular origin and persisting for >24 hours, confirmed by CT scan and clinical examination at baseline. Exclusion criteria included: i) patients who had unconsciousness, aphasia, or severe cognitive impairment that could not cooperate with the verbal clinical interview; ii) patients who had a hemorrhagic stroke, transient ischemic attack, autoimmune diseases, renal or liver failure, or cancer; iii) patients who had psychiatric DSM-5 diagnoses except major depression, bipolar disorder, and dysthymia or were taking antidepressants or other psychoactive drugs; and iv) patients who failed to perform MRI scans for various reasons.

At the time of hospitalization, demographic and vascular risk factors information were collected. Sociodemographic data were collected from semi-structured interviews, including age, gender, religion, years of education, marital status, and occupation. Vascular risk factors identified from underlying diseases included hypertension, diabetes, dyslipidemia, ischemic heart disease, previous stroke, and atrial fibrillation.

Within three months after the onset of stroke, participants were assessed for their depressive symptoms by the Montgomery-Åsberg Depression Rating Scale[31], a 10-item clinician-rated scale. Participants were divided into two groups according to cut-off scores for depression (scores ≥ 7) and non-depression (scores < 7).[32]

### Magnetic Resonance Imaging study

The fluid-attenuated inversion recovery images (FLAIR) from MRI were used to estimate and calculate the location and volume of white matter microstructure hyperintensities with two methods. The first method was manual identification. FLAIR images were adjusted for appropriate contrast and level. The brain images were segmented into four areas via left/right along the longitudinal fissure and frontal/dorsal by central sulcus. The adjusted FLAIR images were detected and measured by a psychiatrist (CJ) and confirmed by a neuroradiologist (NJ) for white matter hyperintensity volume in each segment in cubic millimeters.

We performed a semi-automated WMHs volumetric detection for the second method to compare with the manual WMHs estimation. The imaging data were submitted to Image J application[33]. The contrast value of each patient’s FLAIR image was adjusted to balance the best visibility of WMHs and other tissue. The application calculated the area of WMHs against the background within a brain tissue border. The border was marked by an ellipse shape to determine brain area without the presence of a skull. The area of WMHs was multiplied by the slice thickness of the scanned images, and the volume of WMHs was recorded. The semiautomated method can reduce bias from the manual process. However, an overestimation of WMHs from the inclusion of non-WMHs structures was expected. We also used DWI to calculate volumes of the acute vascular lesion with the same manual method.

### Statistics

The statistical analyses were performed with SPSS 22.0. The study data conformed to a normal distribution, and quantitative data were expressed as mean ± standard deviation. Chisquare test, Cochran’s and Mantel-Haenszel statistics, and Independent-Sample T-test were used for quantitative data. Alpha level of 0.05 or lower was considered statistically significant. Linear regression was used to find the association between depressive symptoms (MADRS score) and volume of WMHs, and volume of the acute lesion from DWI, adjusted with covariates of interest. Association between manual and semi-automated method was examined with a scatter plot and multiple regression.

## Results

### Demographic data of the patients with and without post-stroke depression

The demographic data of the two groups (depressed vs. non-depressed), including age, gender, religion, years of education, marital status, employment status, and underlying diseases, were comparable. A summary of the participants’ demographic data is presented in Table 1. Twenty-nine patients were enrolled, including 15 males and 14 females, aged 55 - 78 years, with an average age of 64.17 ± 6.2 years. Of the 29 patients, 11 (37.93%) met the diagnostic criteria for early-onset PSD. The PSD group (*n* = 11) and non-depressed group (*n* = 18) did not differ in age, gender, religion, marital status, education, occupation, and underlying diseases.

**Table 1.**
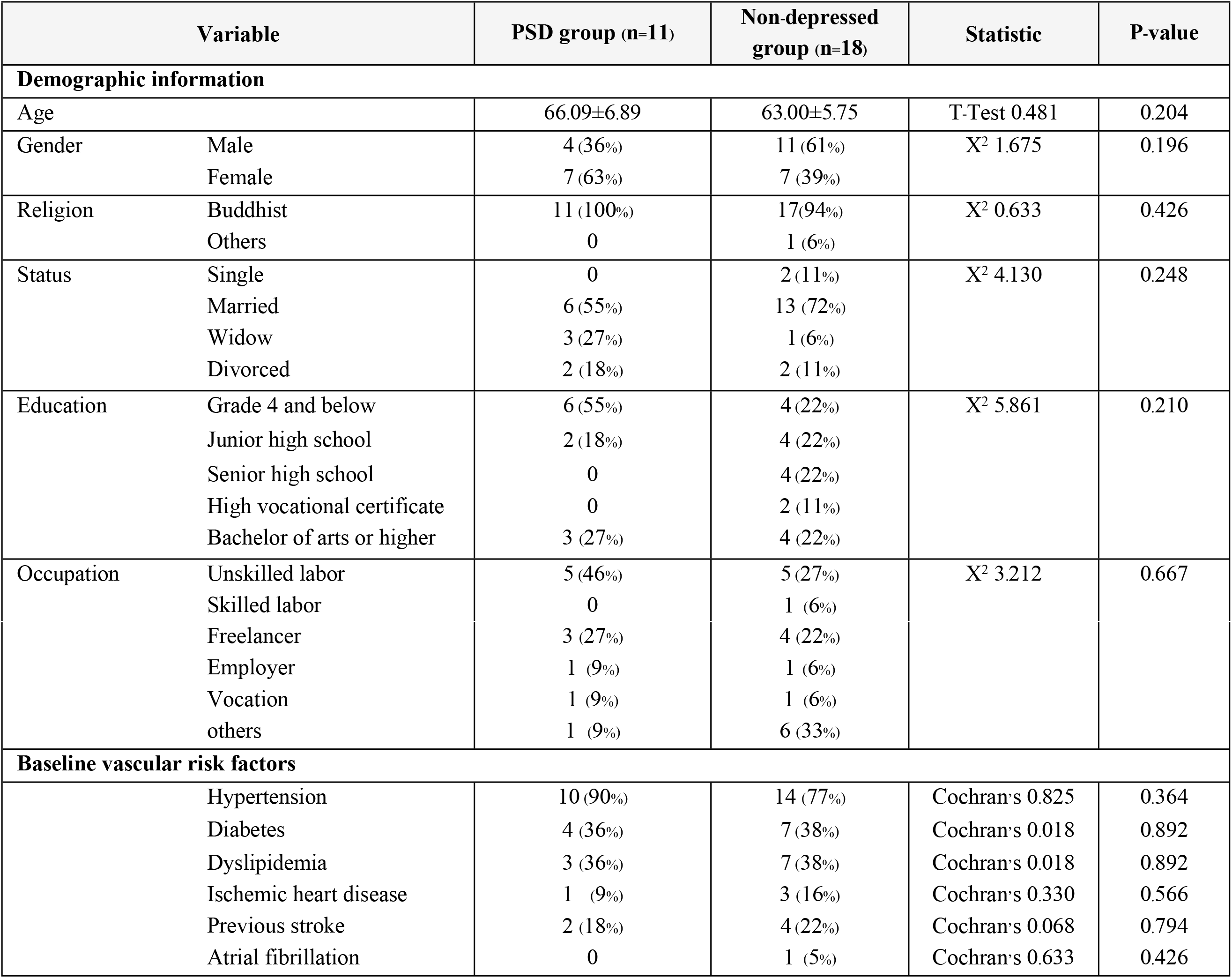
Demographics data and baseline vascular risk factors of participants (n = 29)

### WMHs volume and early-onset post-stroke depression

The mean total volume of WMHs is 17500.97 ± 15145.52 cu.mm. PSD group had the mean total WMHs volume of 17136.45 ± 17500.28 cu.mm. and non-depressed group had the mean total WMHs volume of 17723.73 ± 14054.26 cu.mm.. Total WMHs volume of the two groups was not statistically significant difference. Mean WMHs volume of segments of the brain is presented in Table 2. There were near significant differences between WMHs of the two groups in the left anterior part (t-test 2.058, p=0.055).

**Table 2.** Mean volume of WMHs between PSD group and non-depressed group in each part of brain regions.

| WMHs Regions | PSD group (n=11) | Non-depress group (n=18) | T-test | P-value |
| --- | --- | --- | --- | --- |
| Left anterior part | $3061.86 \pm 790.49$ | $6851.61 \pm 7745.41$ | 2.058 | 0.055 |
| Left posterior part | $6436.16 \pm 14683.37$ | $3118.14 \pm 3657.13$ | -0.923 | 0.364 |
| Right anterior part | $3676.65 \pm 2025.42$ | $4973.08 \pm 2721.13$ | 1.363 | 0.184 |
| Right posterior part | $3961.77 \pm 6550.11$ | $2780.88 \pm 2724.33$ | -0.680 | 0.502 |
| Total WMHs | $17136.45 \pm 17500.28$ | $17723.73 \pm 14054.26$ | 0.100 | 0.921 |

Box plots of WMHs volume in the depressed and non-depressed groups were demonstrated in figure 1. There were four participants in the non-depressed group and two participants in the PSD group who were outliners because they have a large infarction area. In the non-depressed group, participant no. 11 and no. 18 had large areas of single vessel infarction at the left anterior part (45999.60 cu.mm. and 21643.50 cu.mm., respectively). In the PSD group, participant no.29 had a large area of single vessel infarction at the right posterior (23481.40 cu.mm.), and participant no. 13 had a wide area of small vessel infarction at the left posterior part (50554.25 cu.mm.).

**Figure 1.**
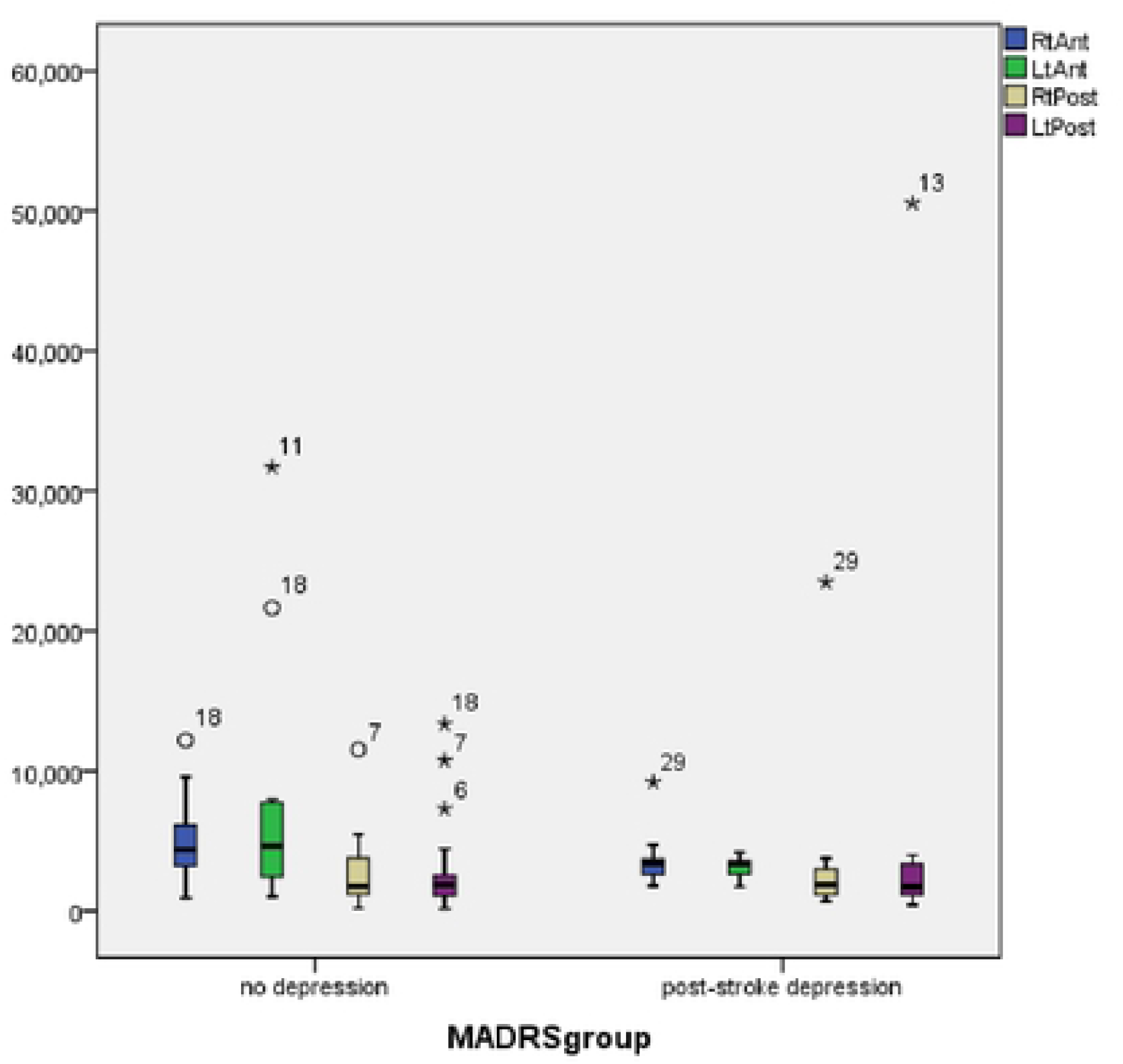
Box plot of volume of WMHs in depress and non-depress group. *Note.* Circles (○) represent outliers within 15-3.0 IQR and stars (*) represent outliers more than 3.0 IQR

### The volume of the acute lesion and early-onset post-stroke depression

The mean total volume of acute stroke lesion measured from DWI is 4945.05 ± 12007.00 cu.mm. The PSD group had the mean total lesion volume of 9276.23 ± 18138.08 cu.mm. and the non-depressed group had the mean total lesion volume of 2298.21 ± 4934.80 cu.mm. The total lesion volume of the two groups was not statistically significant difference. The mean volume of acute lesions segments of the brain is presented in Table 3.

**Table 3.**
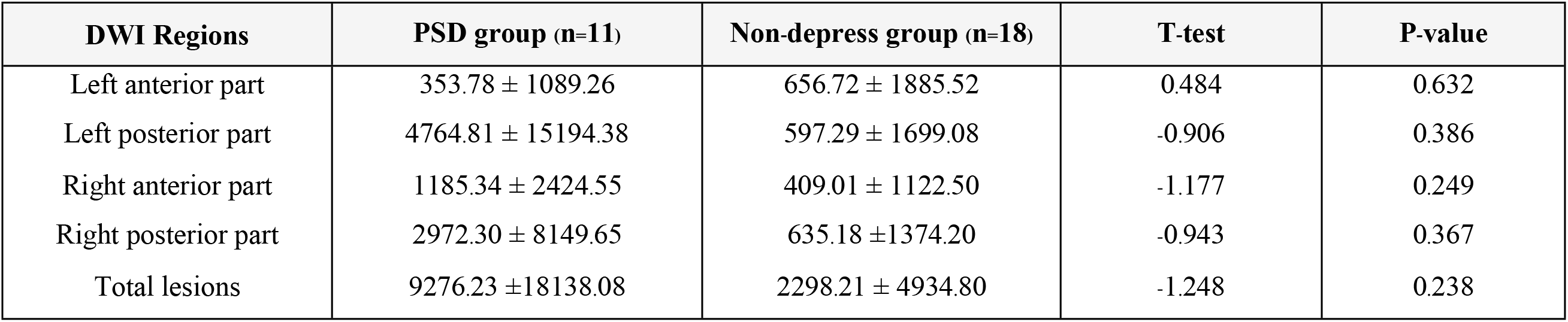
Mean lesions volume from DWI between PSD group and non-depressed group in each part of brain regions.

Figure 2 showed the relationship between Total WMHs volume and MADRS score. The scatter plot showed a dispersion of MADRS score in the lower range of WMHs volume. However, there was no correlation between Total WMHs volume and MADRS score, as evidenced by a linear plot of nearly horizontal alignment.

**Figure 2.**
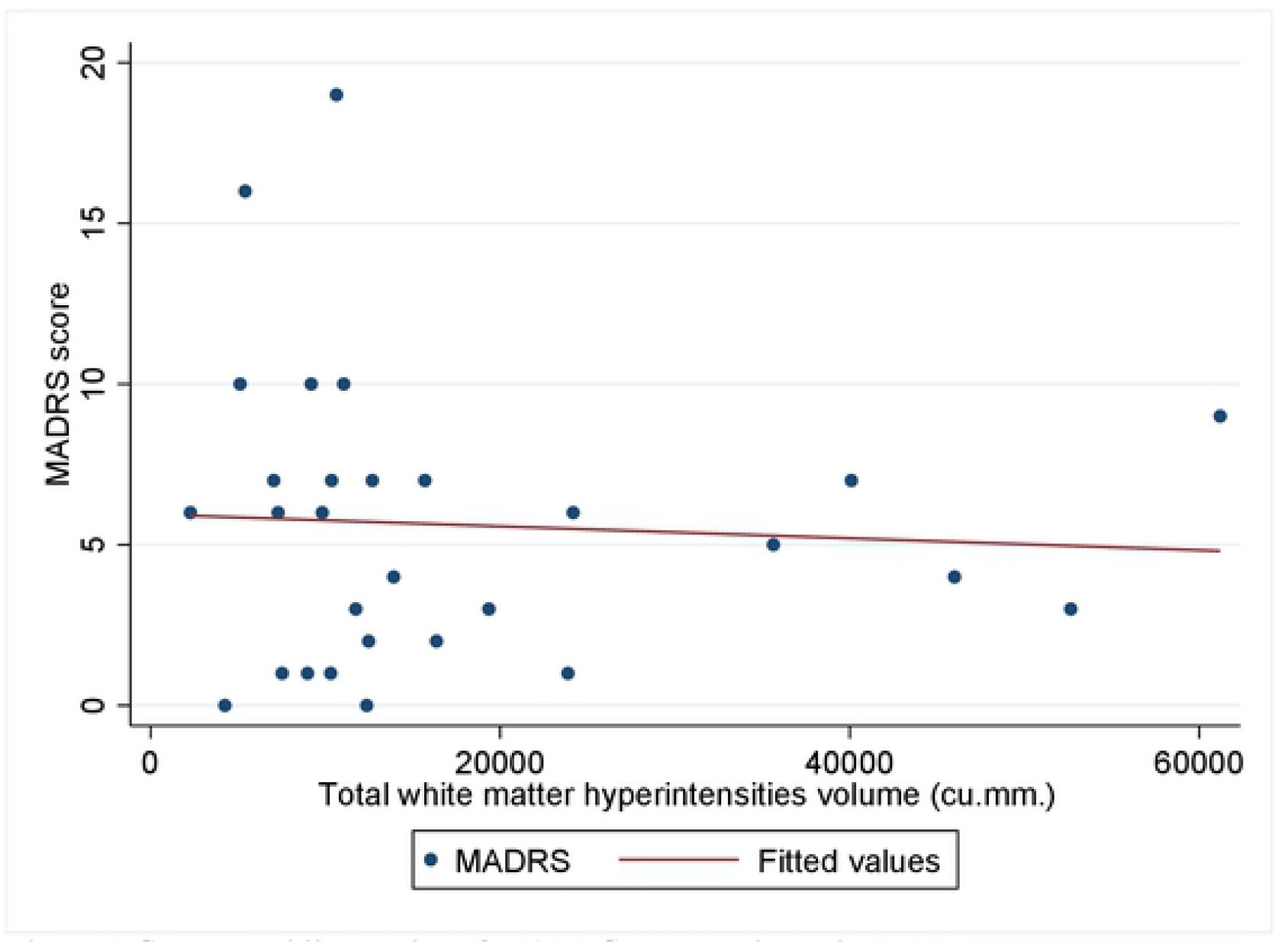
Scatter and linear plot of MADRS scorc and Total WMHs

The results of regression analysis between demographic, clinical, and imaging variables and MADRS score showed in Table 4. In univariate analysis, no variable showed a statistically significant association with MADRS score, including total WMHs and acute lesion volume. Moreover, no association was found between total WMHs and the acute lesion volume and MADRS score in multiple linear regression, adjusting with age, gender, and education level.

**Table 4.** Univariate and multiple linear regression analysis of demographic and clinical variables, including WMHs and volume of acute lesion, and MADRS score. *Note.* Total WMHs and volume of acute lesion were divided by 10,000 before use in the regression analyses.

| Variables | B | SE | p-value | 95% confidence interval |
| --- | --- | --- | --- | --- |
| **Univariate analysis** |  |  |  |  |
| Age | 0.043 | 0.138 | 0.775 | -0.224, 0.326 |
| Female | 2.805 | 1.615 | 0.094 | -0.509, 6.118 |
| Status (married) | 0.795 | 1.784 | 0.659 | -4.454, 2.865 |
| Education (lower than 4 years) | 2.105 | 1.744 | 0.238 | -1.472, 5.682 |
| Occupation (non-labor work) | -0.172 | 1.753 | 0.923 | -3.768, 3.426 |
| Previous stroke | -1.203 | 2.088 | 0.569 | -5.486, 3.081 |
| Total WMHs | -0.186 | 0.571 | 0.747 | -1.357, 0.985 |
| Volume of acute lesion | 0.744 | 0.707 | 0.302 | -0.706, 2.196 |
| **Multiple linear regression analysis** |  |  |  |  |
| Age | -0.066 | 0.166 | 0.696 | -0.409, 0.277 |
| Female | 2.028 | 1.872 | 0.290 | -1.844, 5.900 |
| Education (lower than 4 years) | 2.342 | 2.032 | 0.261 | -1.863, 6.5471 |
| Total WMHs | -0.628 | 0.721 | 0.392 | -2.120, 3.027 |
| Volume of acute lesion | 0.923 | 1.017 | 0.374 | -1.181, 3.027 |

Further analysis of brain regions and MADRS score also showed no association. In addition, logistic regression analysis showed no association between the WMHs and acute lesion volume and depression status (result in Table 5).

**Table 5.** Univariate and multiple logistic regression analysis of demographic and clinical variables, including WMHs and volume of acute lesion, and MADRS score. *Note.* Total WMHs and volume of acute lesion were divided by 10,000 before use in the regression analyses.

| Variables | OR | SE | p-value | 95% confidence interval |
| --- | --- | --- | --- | --- |
| **Univariate analysis** |  |  |  |  |
| Age | -0.066 | 0.166 | 0.696 | -0.409, 0.277 |
| Female | -2.028 | 1.872 | 0.290 | -5.900, 1.844 |
| Education (lower than 4 years) | -2.342 | 2.033 | 0.261 | -6.547, 1.863 |
| Total WMHs | -0.628 | 0.721 | 0.392 | -2.119, 0.862 |
| Volume of acute lesion | 0.923 | 1.017 | 0.374 | -1.181, 3.027 |
| **Multiple logistic regression analysis** |  |  |  |  |
| Age | 1.024 | 0.082 | 0.770 | 0.874, 1.199 |
| Female | 0.605 | 0.591 | 0.607 | 0.089, 4.102 |
| Education (lower than 4 years) | 0.268 | 0.277 | 0.203 | 0.035, 2.032 |
| Total WMHs | 0.548 | 0.311 | 0.290 | 0.181, 1.667 |
| Volume of acute lesion | 2.421 | 1.162 | 0.186 | 0.653, 8.979 |

### Manual versus semi-automated WMHs estimation

The two methods of WMHs volume estimation yielded a similar result. There were 21 pairs of imaging data from the semi-automated method for comparison. The other eight imaging data points were not applicable to the semi-automated method. The manual method reported a mean volume of 18615.74 cu.mm. with a standard deviation of 16198.24 (min-max 5122.3 - 61201.2 cu.mm.). The semi-automated method reported a mean volume of 20986.99 cu.mm. with a standard deviation of 18543.22 (min-max 89.66 - 61604.09 cu.mm.). Pairwise comparison showed a correlation coefficient of 0.75 (p-value < 0.001). A scatter plot shows an estimation in the manual vs. semi-automated method (Figure 3).

**Figure 3.**
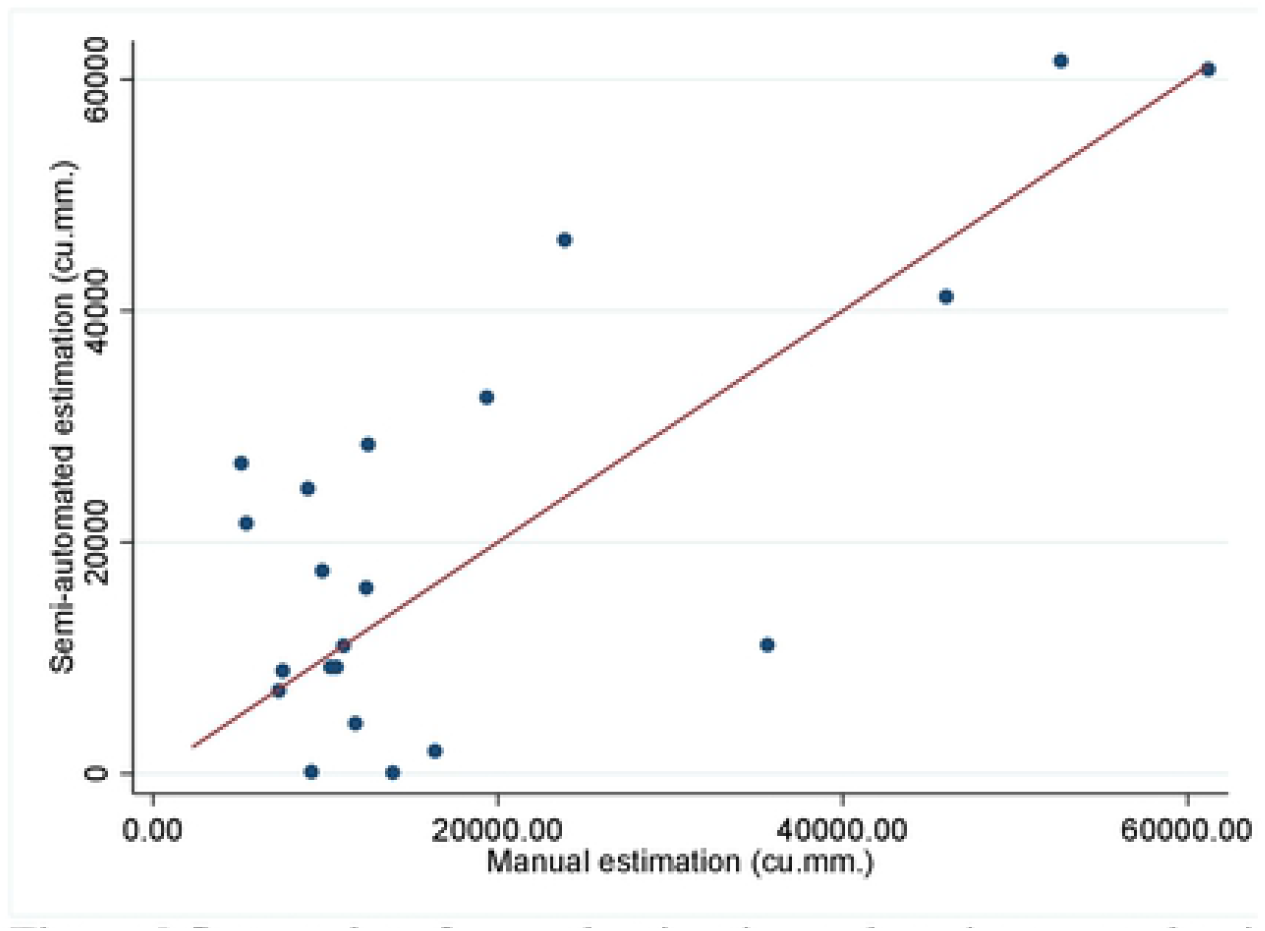
Scatter plot of manual estimation and semi-automated estimation of WMHs. *Note.* Red line shows a reference line of slope = 1.

## Discussion

In the present study, we could not detect a significant association between WMHs volume or volume of acute lesions and PSD. However, previous studies found lesion locations of stroke associated with PSD.[17-24] Our study result may be explained by the neurological track mechanism.[34] The track hypothesis holds that neurotransmitters of noradrenergic neurons and serotonergic neurons travel along the brain’s deep cortex to form a subcortical circuit, including the limbic-cortical-striatal-pallidal-thalamic circuit (LCSPT) [14, 35]. The circuit was correlated with PSD. In the current study, the identified WMHs and the acute lesions might not completely block the LCSPT circuit.

Moreover, the onset of depression following a stroke (early vs. late) should be taken into account. Consistent with this, previous data have shown that no correlation between brain lesions and early onset PSD. Nys et al. (2005) found no association between severity of depressive symptoms and lesion location, presence of preexistent lesions (white matter lesions and silent infarcts), and demographic factors (age, education, and gender) within three weeks after a first-ever symptomatic stroke.[36] Vataja et al. (2001) also concluded that the extent of white matter lesions and atrophy did not differ in patients with and without depression occurring 3-4 months after stroke.[37]

Furthermore, it could be explained from the cytokine point of view that pro-inflammatory cytokines such as CRP and IL-6 are found numerously in patients with stroke with WMHs, and PSD may cause further neuronal damage and increase the later formation of WMHs.[38] Multiple processes have occurred after a stroke event; switched on the release of inflammatory mediators, triggered many processes to neuronal cell death, monocyte infiltrated into the injured vessel wall, activated microglia leading to oligodendrocyte death, followed by degenerated of myelinated fibers that could be seen as white matter damage.[39] Therefore, after a stroke, the formation of WMHs could be seen more in later MRI after a few months, resulting in higher depressive severity in the late-onset type.

Another explanation is previous studies of PSD and WMHs use the Fazekas scale that may be confounded by the physician’s decision. These scales use estimating by visual rating from an overall brain imaging section, not from calculation to total volume. Compared with the volumetric method, the Fazekas scale had lower reliability, lower sensitivity, and lower objectivity.[13] The less accurate estimation of the visual rating scale might be the cause of the false interpretation. Our study detected and measured WMHs volume with more objectivity, and the results should be more accurate. This study also used the semi-automated method to estimate WMHs volume compared with manual estimation and found that they were comparable. Therefore, the result of WMHs volume in the current study could closely represent the brain’s real WMHs lesion. The non-association between WMHs volume and depression might be confirmed.

In addition, the findings suggested that there was a trend of association of WMHs volume in the left anterior segment and depression, which was consistent with previous studies. Robinson et al. [25] found that the severity of depression was significantly increased in patients with left anterior lesions. In contrast, the present research found that demographic data did not show any significant difference between the two groups. Tang et al. [40] also found there was no difference in PSD in age, sex, and previous stroke. Shen et al. [27] showed the same result that gender, age, marriage, and education level were not different between the two groups. However, Shi Y et al. [41] found that female gender, age, family history, the severity of the stroke, and level of handicap were risk factors for PSD.

### Limitations

Results of the current study should be interpreted considering its limitation. First, this study has a cross-sectional design with a small sample size, which limited the statistical power to detect the two groups’ differences. Due to the lockdown measure for the COVID-19 situation in Thailand, MRI was preserved only for stroke patients with an indefinite diagnosis. Second, the current study separated the brain into four areas and cannot reliably determine the underlying tracts within regions. This partitioning limited the ability to look at specific regions beyond gross delineation. Therefore, area of the identified white matter damage might not represent damage in the neural tract. Third, we cannot separate the LCSPT circuit correlated with PSD by using routine MRI protocols which yield the low-resolution, low-angle images. Fourth, some of the MRI examinations were performed after the acute phase of ischemic stroke. Hence post-stroke depression may not present in the acute phase of stroke.[42] To better explore WMH and PSD’s relationship, a follow-up MRI scan later than three months might be suitable. Fifth, some of the DWI examinations were performed within three months after the acute phase of ischemic stroke; hence some ischemic lesions on DWI may disappear.

### Implications and future research direction

The current study showed the possibility of using objective lesion measurements in clinical practice, which required less radiologists’ expertise and therefore were more accessible for other disciplines. However, volumetric measurement of lesions might not be sufficient to predict PSD. Two possibilities were suggested for imaging biomarker: the use of brain imaging technique (e.g., deep tensor imaging [43]) could be applied to specify lesion within the related track; and a data-driven process using machine learning might reveal additional imaging features that related to PSD (radiomic [44]). In addition, various inflammatory and genetic markers have been associated with PSD. The need for further investigation in biological predictors of the early-onset and late-onset PSD is warranted.

### Conclusions

In conclusion, this study’s results suggest that WHMs volume might not contribute to the development of early-onset PSD in older patients from both manual estimation and semiautomated methods. However, WHMs volume of the left anterior segment showed a trend of difference between the two groups. This study showed a possibility of quantitative MRI analysis in clinical practice. However, further investigation in a larger group of patients with a longer post-stroke period is needed.

## Acknowledgements

This study was supported by Rachadapisaek Sompote Fund number 63/073, Chulalongkorn University. The authors appreciate all participants and caregivers for their cooperation, and the Department of Psychiatry, Department of Neurology, Department of Radiology, Faculty of Medicine, Chulalongkorn University, Thailand, for their assistance. The authors appreciated Ms. Kanokvalee Ponkanist for her support with the extraction of brain imaging data.

## Supplementary material

The dataset file for open access will be available in the electronic version of this article.

